# Spatial Transcriptomics Reveals Alterations in Perivascular Macrophage Lipid Metabolism: Insights into the onset of Wooden Breast Myopathy in Broiler Chickens

**DOI:** 10.1101/2023.08.19.553989

**Authors:** Ziqing Wang, Paul Khondowe, Erin Brannick, Behnam Abasht

## Abstract

This study aims to use spatial transcriptomics to characterize the cell-type-specific expression profile associated with the microscopic features observed in Wooden Breast myopathy. 1 cm^3^ muscle sample was dissected from the cranial part of the right pectoralis major muscle from three randomly sampled broiler chickens at 23 days post-hatch and processed with Visium Spatial Gene Expression kits (10X Genomics), followed by high-resolution imaging and sequencing on the Illumina Nextseq 2000 system. WB classification was based on histopathologic features identified. Sequence reads were aligned to the chicken reference genome (Galgal6) and mapped to histological images. Unsupervised K-means clustering and Seurat integrative analysis differentiated histologic features and their specific gene expression pattern, including lipid laden macrophages (LLM), unaffected myofibers, myositis and vasculature. In particular, LLM exhibited reprogramming of lipid metabolism with up-regulated lipid transporters and genes in peroxisome proliferator-activated receptors pathway, possibly through CD36-mediated signaling. Moreover, overexpression of fatty acid binding protein 5 could enhance fatty acid uptake in adjacent veins. In myositic regions, increased expression of cathepsins may play a role in muscle homeostasis and repair by mediating lysosomal activity and apoptosis. A better knowledge of different cell-type interactions at early stages of WB is essential in developing a comprehensive understanding.

## Introduction

Wooden Breast myopathy (WB) has been a major cause for economic loss in the poultry industry due to deterioration to the muscle integrity and meat quality, rendering pale and stiff p. major muscle in the cranial to caudal regions [1]. High-throughput sequencing and omics techniques have previously revealed various factors and related expression patterns associated with WB such as energy metabolism, hypoxia, oxidative stress and cellular signaling [2–5].

Nevertheless, skeletal muscle is a complex tissue where multiple cell populations reside and work in conjunction, from myofibers, vasculature, nerve fibers, and connective tissues including collagen producing fibrocytes, adipocytes, and immune cells. While single-cell RNA-seq surpasses whole transcriptome RNA-seq in yielding expression profiles at cell type level, the spatial relationships of interrelated cells within the tissue are lost upon processing. The advent of spatial transcriptomics has introduced a novel approach allowing mapping omics changes to individual architectural regions and structures and at times to individual cell populations histologically within tissue. For example, spatial transcriptomics techniques have been used to identify a plaque-induced gene network which constitutes a coordinated cellular response between astrocytes and microglia in the vicinity of amyloid plaques in Alzheimer’s Disease [6]. Furthermore, transcriptional signatures of functional anatomical domains were characterized in both murine muscle tissue after nerve crush injury [7] and fish myotomal muscle [59]. Spatial analysis using a rabbit model of rotator cuff tear revealed heterogeneous myofibril lesions and identified markers for regenerating myofibers [60]. To the best of our knowledge, there are no published research performing spatial transcriptomics on chicken muscles, either with or without myopathic changes.

WB disease shows unique pathological features, starting with lymphoplasmacytic phlebitis progressing to complete venous occlusion with sparing of adjacent arteries [24]. Additionally, veins in myopathic regions frequently are lined or cuffed by clusters of lipid-laden macrophages. In degenerated myofiber regions, segmental disruption, fiber splitting and vacuolation are followed by fibrosis, or scarring, alongside multifocal myoregeneration, characterized by myotube formation with nuclear rowing [24]. At present, it is uncertain which cells or tissue regions within p. major muscle are responsible for the omics changes identified. By elucidating molecular changes localized to certain cells or tissue structures associated with histological features, we can further characterize the pathophysiology and pathogenesis of WB. Furthermore, this investigation may provide valuable insights for diagnosis, treatment or prevention. Spatial transcriptomics is applied in order to overlay transcriptome on top of histology for the purpose of characterizing expression pattern particular to resident cell types and their potential interactions. Consequently, this study aimed to improve our understanding on functional roles played by distinct cell types and tissue architectural elements during WB development.

## Results

### Sample classification and Gene expression metrics

The stitched high-resolution images of H&E-stained samples are shown in Fig. 1A. A more compact muscle tissue was observed in Sample 1 and 2 than in sample 3. Furthermore, sample 3 exhibited notable pathological lesions associated with WB, including myositis, evident as infiltration of immune cells in the interstitium (Fig. 1A). Accordingly, sample 3 was classified as affected in contrast to sample 1 and 2 as unaffected. The presence of lipid laden macrophages (LLM) alone was not accompanied with nor an accurate indicator of myopathic changes as muscles both with and without myositis demonstrated small to moderate numbers of LLM accumulating around veins. Nevertheless, the presence of LLM in these samples could imply a certain degree of white striping and cellular changes, potentially culminating in the development of WB beyond the age at which samples were collected. In terms of histopathologic analysis, feature appearance ratio was calculated as number of tiles containing that feature divided by total number of tiles occupied by the sample. As a result, ratio of LLM increased from 8% for unaffected sample 1 and 2 to 27% and 42% for affected sample 3_1 and 3_2, respectively. Accumulation of LLM was observed adjacent to or circumferentially cuffing veins in both affected and unaffected samples (Fig. 2B), with noticeably increased number and severity in affected tissue. Similarly, area of connective tissue between myofiber bundles was seemingly larger with higher ratio in the WB affected samples (Fig. 1A). Focal myositis was only observed in the affected muscle and manifested a sporadic pattern. The ratio of myositis increased from 20% in sample 3_1 to 60% in 3_2, corroborating that WB lesions happen sporadically, often in a multifocal polyphasic manner, and affect myofibers at different depth within the breast muscle.

**Figure 1.**
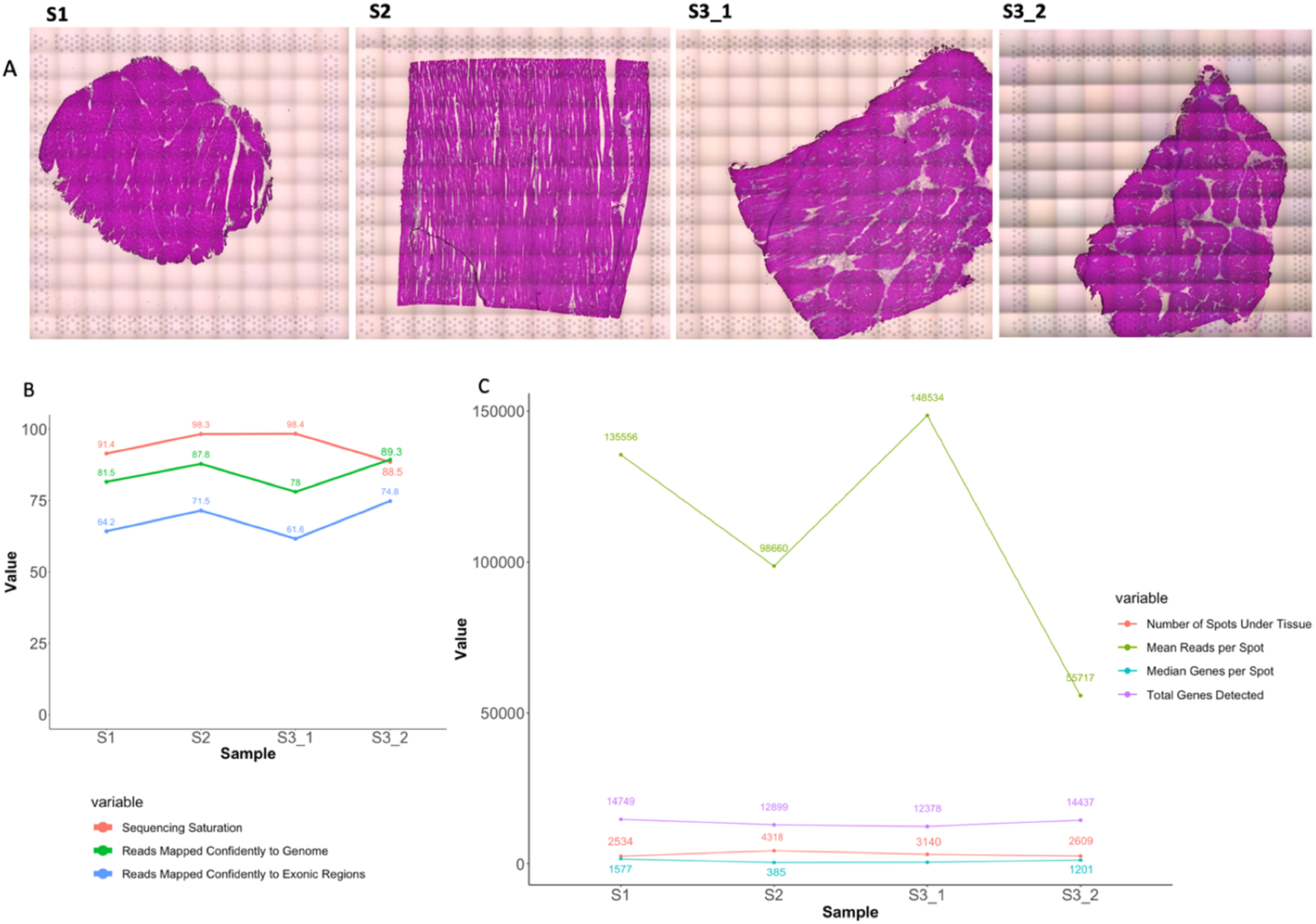
**A)** Stitched images of Hematoxylin and eosin (H&E) stained pectoralis major samples from broiler chickens on the Visium gene expression slides (10xGenomics) for spatial transcriptomics. The grid pattern indicates tile borders. **B)** Sequencing saturation and mapping percentage of reads from broiler chicken pectoralis major muscle mapped confidently to the chicken genome. **C)** Statistics of number of reads, spots and genes per spot. S1: sample 1 (oblique); S2: sample 2 (longitudinal); S3_1: sample 3 processed with permeabilization time of 18 minutes (oblique); S3_2: sample 3 processed with permeabilization time of 6 minutes (oblique). S1 and S2 samples were classified as “Unaffected” and Samples S3_1 and S3_2 were classified as “Wooden Breast affected” by routine histologic analysis.

**Figure 2.**
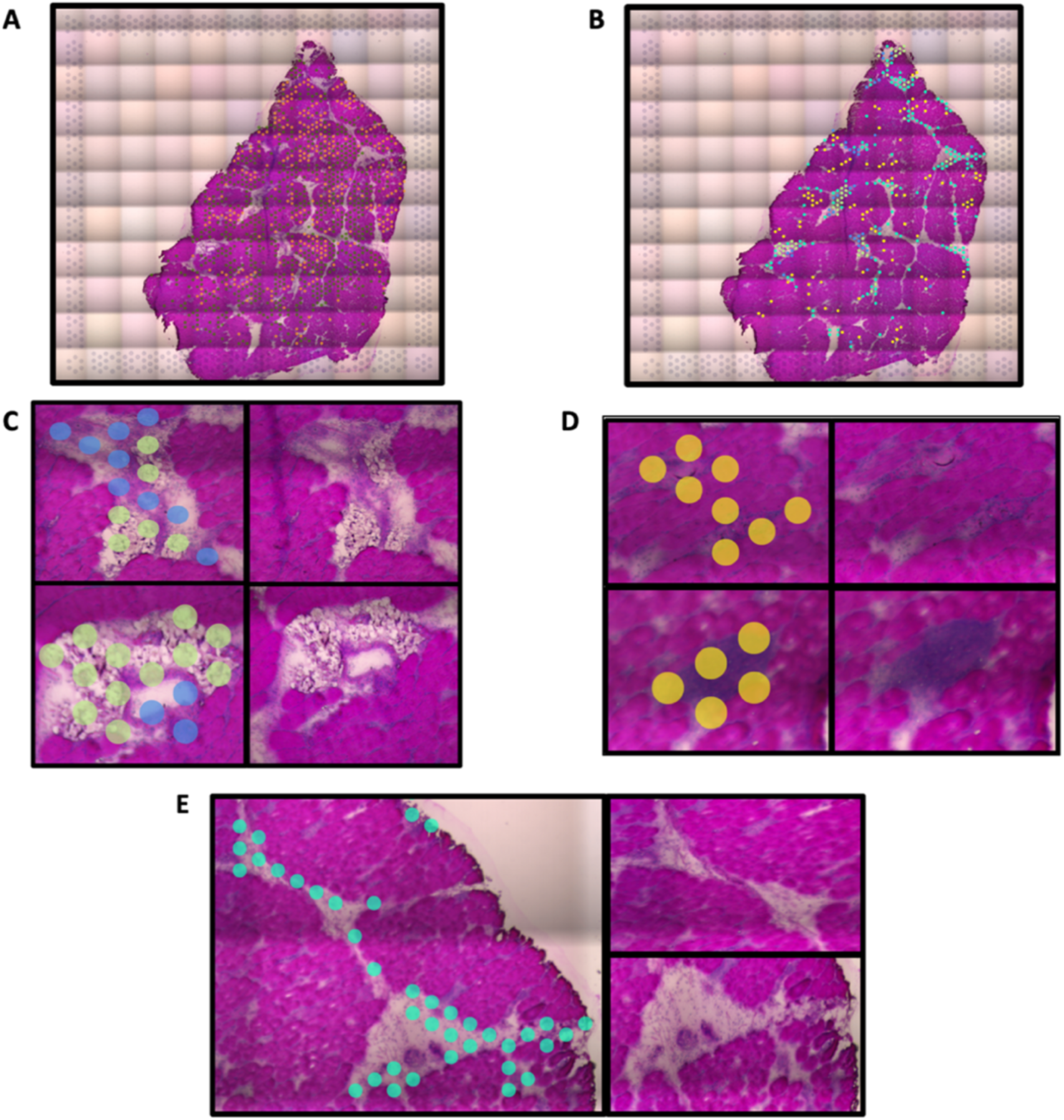
Representative spatial transcriptomics cell type clusters determined by K-means clustering. Clustering is identified by the presence of colored dots. Unmarked insets to the right of each region demonstrate the microscopic appearance of the cells/tissue structures in each region for visual histologic confirmation of cluster classification by cell type or feature. **A)** Myofibers (orange and dark green dots) clustered separately from **B)** Non-myofiber features (no dots). At higher magnification, clustering separated **C)** vessels (blue dots) from perivascular lipid-laden macrophages (light green dots), **D)** inflammatory infiltrates indicative of myositis (yellow dots), and E) perimysial connective tissue (teal blue dots).

Sequencing and mapping performance measurements were satisfactory and comparable among samples (Fig. 1B and Fig. S1). Despite a relatively lower sequencing depth, percentage of reads mapped to the genome and exons, as well as number of genes detected increased for sample 3 after changing permeabilization time to 6 minutes (Fig. 1B and 1C). Overall, these summary metrics suggested an adequate experiment for downstream feature classification.

### Unsupervised K-means clustering

Unsupervised K-means clustering identified 5 different unique histologic features, cell types, or regions, including LLM, myositis, vessels (specifically veins), connective tissue, and myofibers (Fig. 2). Myofibers manifested heterogeneity in expression profiles, as identified by multiple clusters (marked by the orange and green dots in sample 3_2 in Fig. 2A) overlaying visually confirmed myofiber clusters. While K-means clustering of Visium spots identified LLM in all the samples, vascular regions were identified only in sample 1 and 3_2 by this analysis. Vessel walls were confirmed microscopically based upon observation of the endothelial-lined lumen, smooth muscle-lined vascular walls, and intraluminal red blood cells, when present. Connective tissue was also identified by K-means clustering in all the samples and confirmed by observation of discrete extracellular collagen fiber bundles between the myofibers or between vessels and muscle fascicles (Fig. 2E). In accordance with histopathologic analysis, myositis was unique to affected sample 3 (3_1 and 3_2) (Fig. 2D), in which distorted and smaller myofibers were infiltrated by immune cells such as histiocytes and heterophils.

Interestingly, some marker genes in one of the myofiber clusters of sample 1 and 2 were vasculature related, such as hemoglobin subunit alpha 1 (*HBA1*) and hemoglobin subunit epsilon 1 (*HBBA*). Therefore, these spots likely indicate the small capillaries abutting myofibers. Albeit virtually invisible histologically, capillary location was suggested by the linear pattern of the spatial transcriptomics spots (Figure 3).

**Figure 3.**
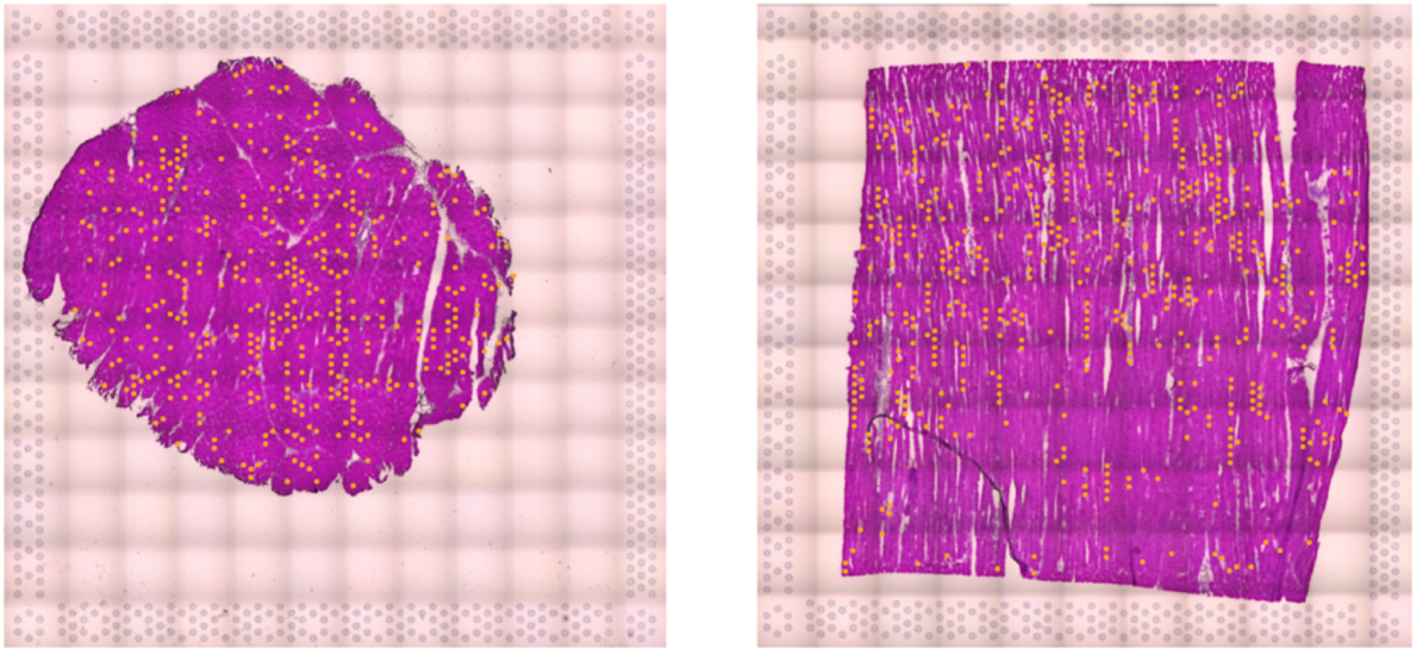
The cluster capturing small capillaries (orange spots). Note the linear arrangement of spots alongside myofibers suggestive of microvascular architecture. **Left:** Sample 1 (oblique); **Right:** Sample 2 (longitudinal).

Marker genes for LLM, connective tissue, myositis and vasculature were reported in Table 1. These genes were selected as marker genes, because (i) they were up-regulated in the clusters identifying these features and (ii) they were overlapped across all 3 samples, except for “vasculature” with overlap in at least 2 samples and myositis in sample 3. Interestingly, markers for LLM were mostly involved in lipid metabolism, indicating their role in processing and storing lipids. In terms of myositis, marker genes encoding lysosomal proteins including cathepsins (*CTSA, CTSK*), IFI30 Lysosomal Thiol Reductase (*IFI30*) and CD74 Molecule (*CD74*) suggested active lysosome and immune cells activities in muscle turnover. As expected, marker genes related to extracellular matrix and smooth muscle were detected by the clusters identifying connective tissue and vascular regions, respectively.

**Table 1.**
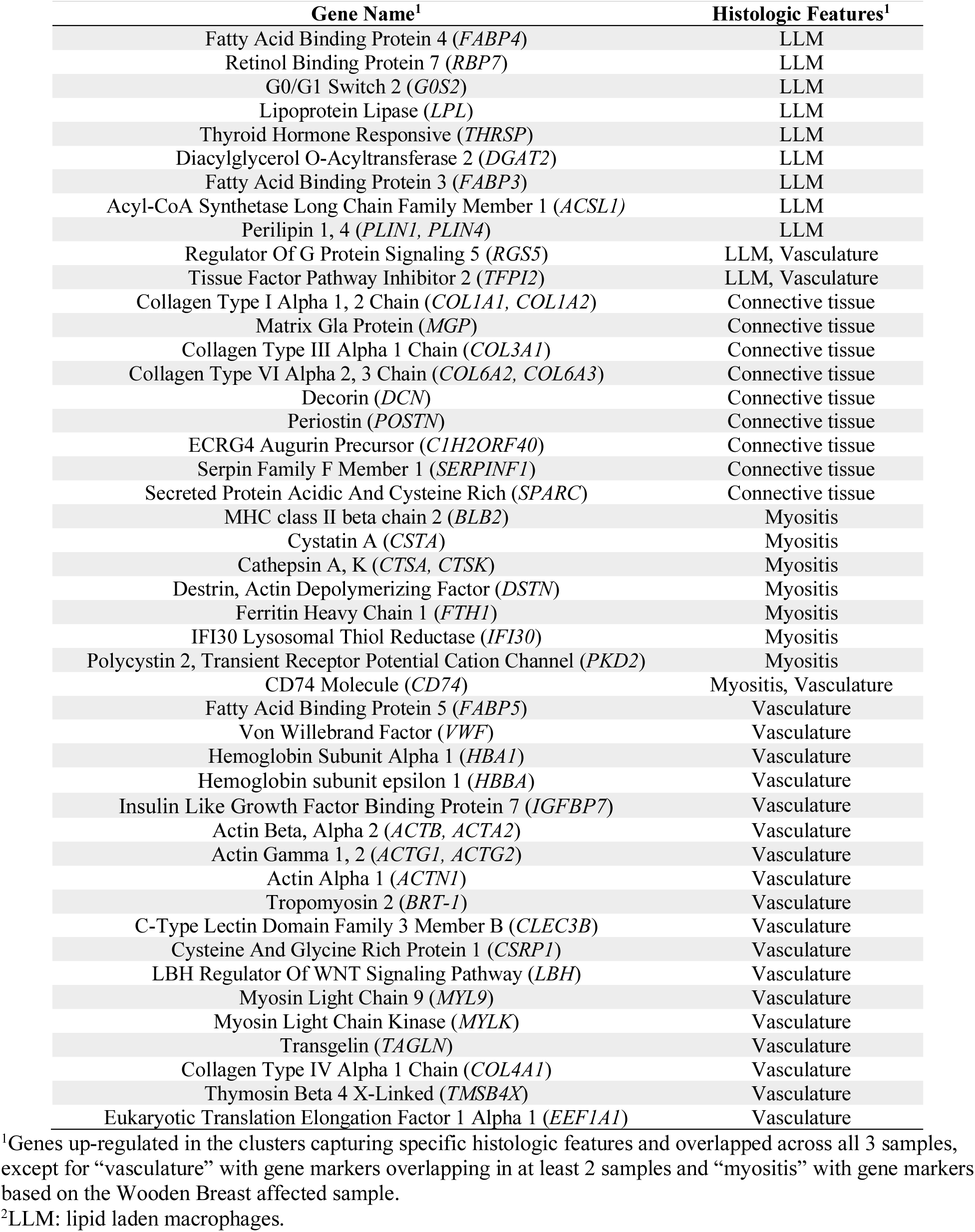
Marker genes for histologic features captured by unsupervised K-means clustering of Visium spots.

### Integration analysis

Integration analysis was performed to identify cell types and marker genes conserved in both WB affected and unaffected samples, as well as to compare between two conditions to find cell-type specific responses to the pathological state. In total, 8 clusters were identified, and UMAP revealed a good overlap of data points between the affected and unaffected conditions (Fig. 4A). Each cluster was then characterized based on their marker genes, resulting in LLM, connective tissue, vascular regions, and myofiber clusters. Identification of 4 distinct clusters characterized as myofibers (M1 to M4) corroborated myofiber heterogeneity found by unsupervised K-means clustering. Heatmaps created using top 10 marker genes in each cluster showed a clear separation of the last 6 clusters (Fig. 4B), with M1 having only 3 markers. Considering the small number of markers in M1 and how scattered the spots in M1 were, this cluster probably contained the spots representative of background information and the ones overlapping across multiple cell types, for instance muscle and connective tissue (Fig. 4A). Moreover, there is a good overlap between these markers and the ones in Table 1, especially for LLM and connective tissue, having 8 and 7 overlapping genes, respectively, further supporting the M1 region as an overlap between myofibers and adjacent tissue.

**Figure 4.**
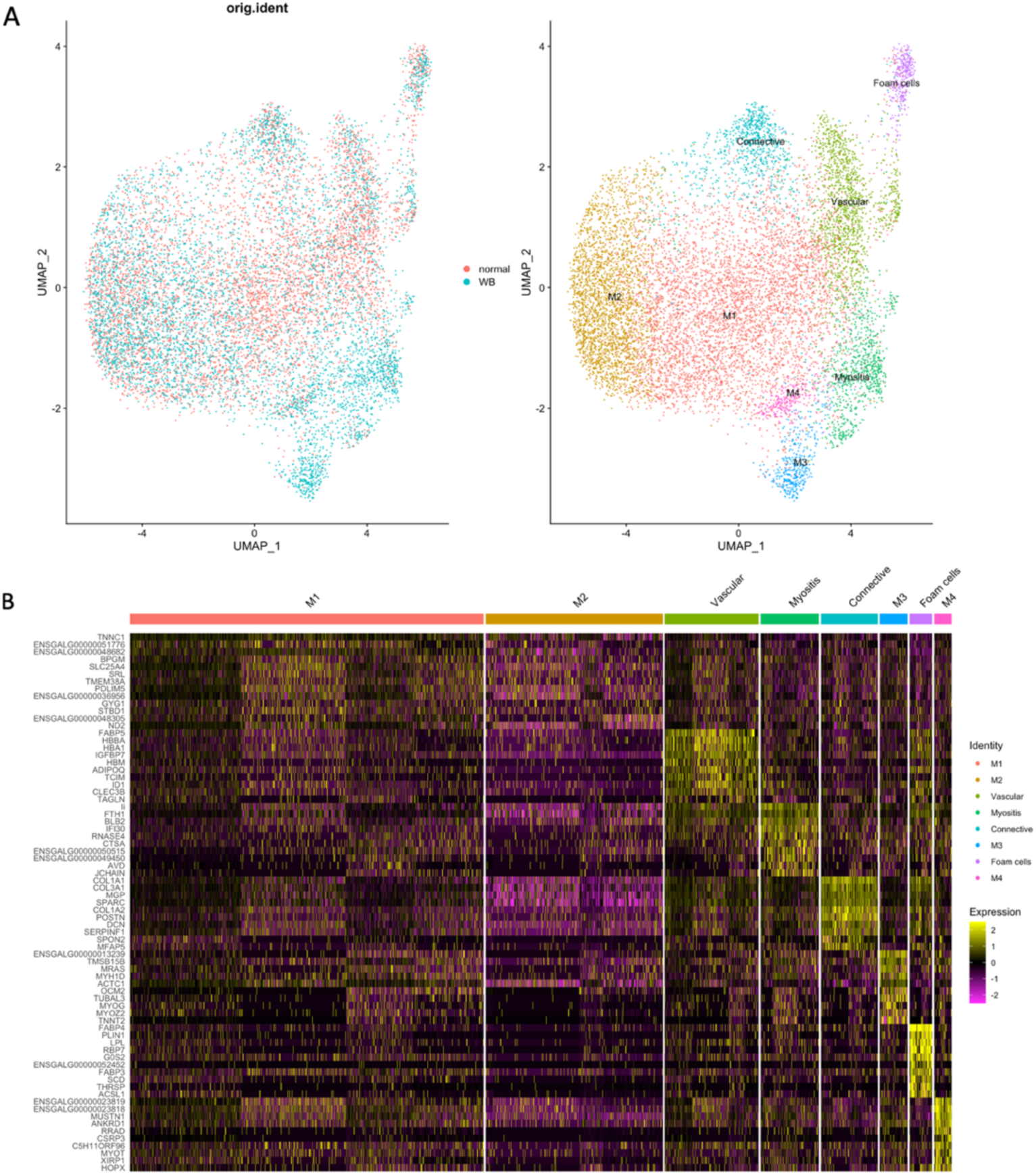
Integration analysis using Seurat. **A)**: Scatter plot of Uniform Manifold Approximation and Projection results. **B)**: Heatmap of the top 10 marker genes of each cluster. In total, 8 clusters were identified by the integration analysis, and then characterized based on their marker genes as lipid leaden macrophages (LLM), connective tissue, vascular regions, and 4 distinct myofiber (M1 to M4) clusters.

In the myopathic condition, connective tissue and vasculature exhibited divergence in expression of some marker genes (Fig. 5). In particular, type I and III collagens (*COL1A1, COL3A1*) showed comparable expression level in both WB affected and unaffected groups, whereas the fibril-associated collagen type XII alpha 1 chain (*COL12A1*) was highly expressed in the myopathic condition (Fig. 5A). Vasculature markers specific to the myopathic condition were EGF like domain multiple 7 (*EGFL7*) and monoamine oxidase A (*MAOA*) (Fig. 5B).

**Figure 5.**
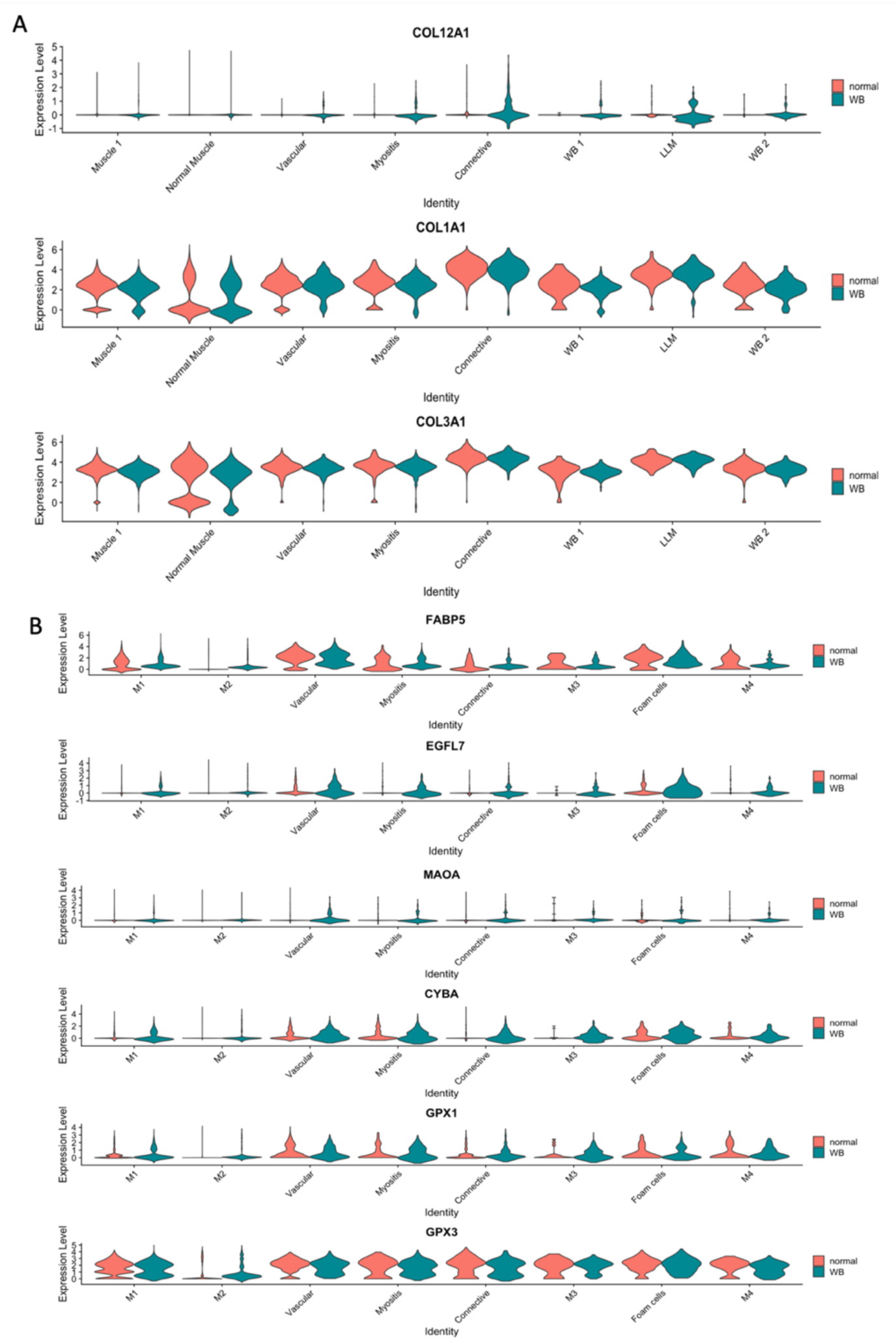
Violin plots of some marker genes in Wooden Breast affected and unaffected samples showed gene expression pattern specific to the myopathic condition. **A)**: Connective tissue and **B)**: Vasculature. These violin plots showed genes expressed specifically to the affected tissue, including collagen type XII alpha 1 chain (*COL12A1*) for connective tissue and EGF like domain multiple 7 (*EGFL7*) as well as monoamine oxidase A (*MAOA*) for vasculature.

Pathway enrichment analysis identified three groups of pathways for LLM, including translation, lipid metabolism and immune response (Fig. S2A). Among pathways for lipid metabolism, the peroxisome proliferator activated receptor (*PPAR*) signaling pathway stood out by sharing 6 top marker genes: fatty acid binding protein (*FABP3, FABP4*), acyl-CoA synthetase long chain family member 1 (*ACSL1*), stearoyl-CoA desaturase (*SCD*), perilipin 1 (*PLIN1*) and lipoprotein lipase (*LPL*). Pathways for connective tissue (Fig. S2B) were mostly related to the extracellular matrix and collagen, suggestive of early tissue remodeling or initiation of fibrosis in WB tissue. As for myositis (Fig. S2C), enriched pathways included lysosome, heterophil degranulation, phagosome and apoptosis consistent with inflammation and clearance of damaged myofiber segments. Top-ranked pathways for myofiber cluster M2 were concentrated on energy metabolism, such as citric acid cycle, oxidative phosphorylation and glucose metabolism (Fig. S3A), whereas those for M4 were focused on cell cycle and apoptosis (Fig. S3B) suggesting possible clean-up/repair. Only three pathways were enriched for M3: 1) infectious disease, 2) rRNA processing and cooperation of Prefoldin and 3) TriC/CCT in actin and tubulin folding. Considering the metabolic pathways and the distance between M2 and M3 to M4 clusters (Fig. 4A), M2 was presumably representing normal myofibers in contrast to damaged or reparative myofibers (M3 and M4) identified in the WB affected sample. As described previously, the M1 cluster genes overlapped with the connective tissue clustering and may demonstrate the myofiber/connective tissue interface.

## Discussion

### Lipid-laden macrophage accumulation

Lipid-laden macrophages (LLM), more colloquially known as “foam cells”, are monocyte/macrophage lineage cells with intracellular accumulation of lipid droplets [14]. This cell population was reported in a previous study to aggregate in perivascular connective tissue in more than 65% of chickens from a purebred commercial broiler line by one week of age [24]. Noticeably, the perivenous (i.e. external perivascular cuffing) location of these LLM in broiler chickens is rather different to that in atherosclerosis where LLM accumulate in subendothelial space and smooth muscle layers within the wall of arteries [47].

*PPARγ* signaling activation in the cluster marking LLM, as indicated by the up-regulation of CD36 molecule (*CD36*), NPC intracellular cholesterol transporter 2 (*NPC2*), *LPL, FABP4* and *FABP3*, suggest similarities between LLM accumulation in chicken breast muscle and metabolic activation of adipose tissue macrophages in human obesity [15]. It is possible that similar metabolic pathways are activated in broiler chickens due to intensive genetic selection for efficient and enhanced muscle growth in commercial meat production, even though the birds do not become obese via lipid deposition in adipocyte depots like their human counterparts. Induction of *PPARγ* activity in adipose tissue macrophages has been previously studied and linked to exposure to elevated levels of oxidized low-density lipoproteins (oxLDL) or palmitate [15]. Accumulation of palmitate and oxLDL were reported in WB [2,4]. Particularly, oxLDLs build-up was indicated by higher levels of its principal components 9-HODE and 13-HODE [17]. Additionally, using a murine alveolar macrophage cell line, Hou and colleagues [18] reported induction of LLM formation along with elevated transcription of *CD36* and *PPARγ* by oxLDL, with significant attenuation by either blocking *CD36* or inhibiting the activity of *PPARγ*. Consequently, LLM formation in broiler chickens likely resulted from a similar process involving oxLDL-*CD36* mediated signaling and *PPARγ* induced up-regulation of its transcriptional target genes [16–18]. A comparable LLM activation in these aforementioned varied tissue origins shared one similarity, a lipid rich extracellular environment.

Figure 6 depicts our proposed mechanism for LLM formation within the p. major muscle of broiler chickens. Basically, *LPL* preferentially hydrolyzes triacylglycerols (TAG) in extracellular space [21]. Upon oxLDL stimulation, scavenger receptors *CD36* and *NPC2* promote lipid uptake into the cells [48]. Specifically, oxLDL binds to its high-affinity receptor *CD36* to enter the cells by endocytosis [52]. *PPARγ* could be activated by both oxLDL and fatty acids (FA), which in turn increase FA uptake by inducing expression of *FABP4* and *CD36* [19]. Once inside, *FABP4* then facilitates FA transport to intracellular vesicles and organelles to boost cholesterol ester and TAG accumulation [20], participated by enzymes stearoyl-CoA desaturase (*SCD*) [49] and diacylglycerol O-acyltransferase 2 (*DGAT2*) [50] respectively. Expression of *SCD* and apolipoprotein 1 (*APOA1*), indicators of elevated cholesterol efflux and biosynthesis of mono unsaturated FA [47], coincided with the significantly elevated lipid level in WB [2, 4, 69]. Unlike in adipose tissue of high feed efficiency broilers (low-fat phenotype), G0/G1 switch gene 2 (*G0S2*) was activated in the LLM, shifting towards a metabolism favoring lipid deposition [25]. Moreover, oxLDL-*CD36* mediated signaling is associated with mitochondrial accumulation of long chain FA, possibly a consequence of enhanced FA oxidation and/or biosynthesis channeled by *ACSL1*, and suppression of LLM migration which could explain the perivascular accumulation and cuffing by these cells [16,22]. Finally, as a constitutive protein of lipid droplets, up-regulation of perilipins (*PLIN1, PLIN4*) was directly linked to increase in lipid droplets [23]. Interestingly, these lipid genes *FABP4, G0S2* and *PLIN1* showed a significantly higher expression in the cranial area than the caudal region [68], suggesting a higher degree of lipid metabolism at play and consistent with the enhanced vulnerability to WB in the cranial region of the p. major muscle.

**Figure 6.**
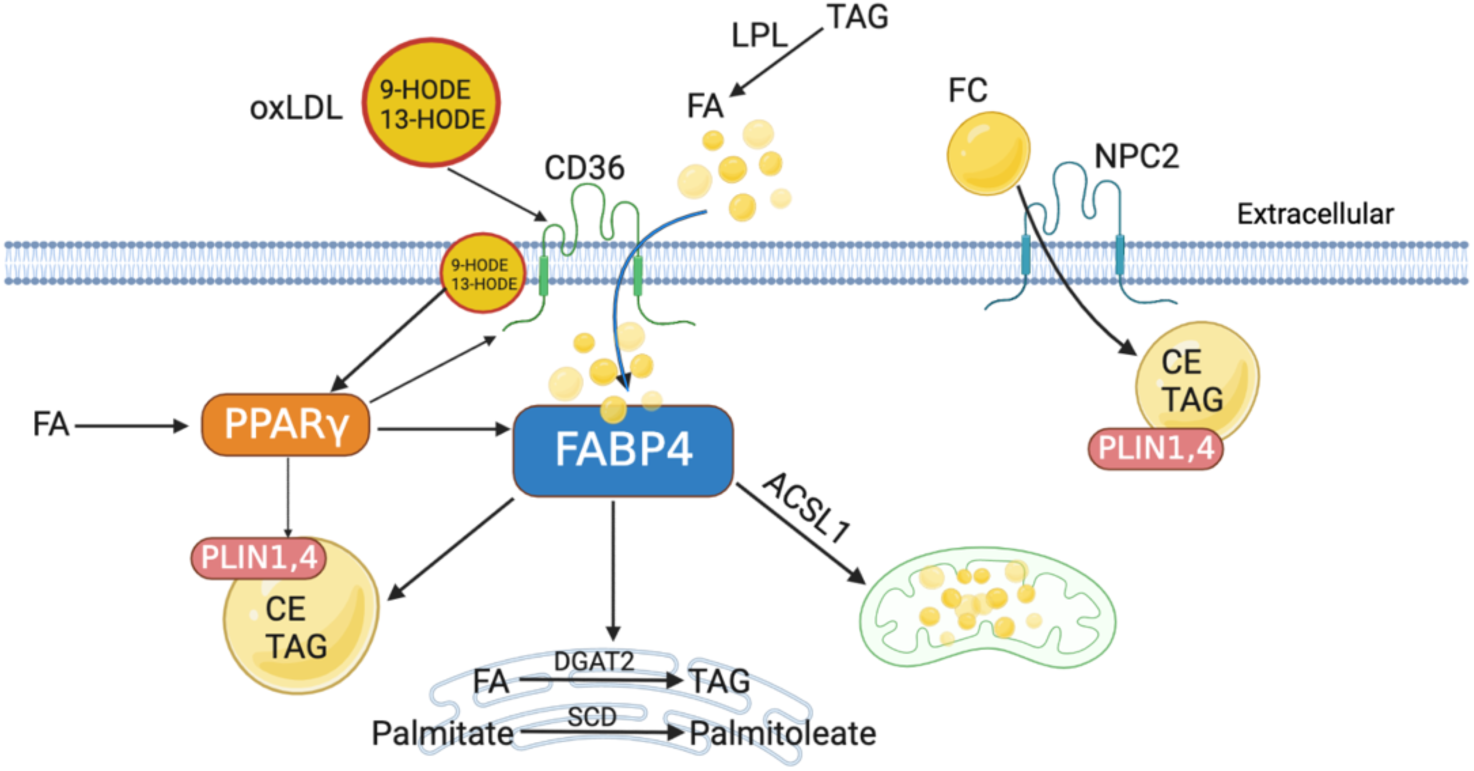
Proposed mechanism of lipid laden macrophage accumulation in the p. major muscle of broiler chickens. Cells are metabolically stimulated by oxidized low-density lipoproteins to activate PPARγ signaling pathway, which boosts lipid uptake and synthesis intracellularly to process excessive surrounding lipids. [oxLDL: oxidized low-density lipoproteins; FA: fatty acids; TAG: triacylglycerol; FC: free cholesterol; CE: cholesterol ester; *CD36*: CD36 molecule; *NPC2*: NPC intracellular cholesterol transporter 2; *FABP4*: Fatty Acid Binding Protein 4; *PPARγ*: peroxisome proliferator activated receptor gamma; *PLIN1,4*: Perilipin 1, 4; *DGAT2*: diacylglycerol O-acyltransferase 2; *SCD*: stearoyl-CoA desaturase; *ACSL1*: acyl-CoA synthetase long chain family member 1]

In general, these perivascular LLM in chicken breast muscle showed comparable gene expression pattern to atherosclerotic LLM, except for genes pertinent to pro-inflammatory cytokines nor M1 and M2 macrophages markers [53,58]. Previous research [16,20,22] on peritoneal and myeloid monocyte-derived LLM in mice and *in vitro* cell culture studies discovered inflammatory activities producing interleukins, tumor necrosis factor (*TNF-α*), monocyte chemoattractant protein (*MCP-1*), etc. Nevertheless, according to a study by Kim et al. [14], non-foamy macrophages are responsible for expressing interleukins and other inflammatory cytokines in the atherosclerotic intima. Since the identified marker genes were primarily related to a lipid metabolism, this LLM population in broiler chickens may primarily aid in processing excessive lipids entering the breast muscle mediated by *LPL*. Alternatively, given the elevated expression of *LPL* in LLM, it is possible that this cell population contributes to perivascular lipid infiltration, thereby establishing a positive feedback loop (i.e. further increasing perivascular lipid accumulation and perivascular LLM attraction) and playing a significant role in the development of the observed pathology. Up to 50% reduction of atherosclerotic lesions was reached in mice deficient for macrophage *LPL* [51]. Furthermore, considering its adjacent location to veins, continuous accumulation of LLM could compress the space occupied by veins or alter vascular flow, eventually leading to phlebitis as seen in older broiler chickens with WB [24].

### Connective tissue and Veins

Myofibers are surrounded by and embedded in connective tissue, or extracellular matrix (ECM), composed of netted collagen fibers and other macromolecules [35]. As a signaling hub, the connective tissue modulates muscle growth, repair and contractile force transmission [35], and thus is vital for muscle integrity. Naturally, the fibril-forming *COL1A1, COL1A2, COL3A1* as well as microfibrillar collagen VI (*COL6A2, COL6A3*) were the conserved markers for connective tissue in this study. The unique expression of *COL12A1* in the myopathic condition is consistent with prior bulk RNA-seq studies showing an up-regulation this gene in WB affected chickens comparing to their unaffected counterparts starting as early as week 2 [5, 65]. Since collagen XII binds the macromolecules in ECM to form bridges that organize adjacent collagen fibrils [36], its deposition indicated a divergence in the structure and biomechanical properties of ECM between WB affected and unaffected tissues. An intense collagen XII deposition was previously reported in regenerative heart of Zebrafish, with involvement in tissue cohesion and regeneration [38]. In correspondence with the stiff characteristic of WB muscle, *COL12A1* expression may be related to constant tensile strain and fibrosis [36]. In human and mice, mutations in *COL12A1* led to joint hyperlaxity and muscle weakness due to a passive force-transducing elastic element in ECM [37]. The enriched pathways of vascular marker genes such as leukocyte transendothelial migration suggested the presence of inflammation in the vasculature within the p. major muscle (Fig. S2D). Although examination of histological images for this study revealed no microscopically visible lesions of phlebitis, expression of some markers certainly imposed risks for venous inflammation in a later myopathic stage (Fig. 5B).One potentially important biomarker was von Willebrand factor (*VWF*), which mediates vascular inflammation via recruitment of immune cells including macrophages, platelets and leukocytes [39]. Furthermore, capillary endothelial *FABP5* could play a role in boosting FA uptake through endothelial walls [19-Iso,2013], thus originating the enhanced lipid deposition in breast muscle of broilers. *FABP5* may also stimulate cell proliferation by regulating *PPARδ* signaling [40]. A rather specific expression of *EGFL7* in the myopathic condition (Fig. 5B) suggested hypoxia-induced angiogenic behavior or vessel regeneration [41]. In addition, adiponectin (*ADIPOQ*) expression in veins was found relevant to peroxidation substrates in the vascular wall [42]. Oxidative stress in the veins of WB affected muscle was inferred by marker genes involved in oxidative stress response pathway, *MAOA*, glutathione peroxidase (*GPX1, GPX3*) and cytochrome B-245 alpha chain (*CYBA*). While *MAOA* and *CYBA* are responsible for production of reactive oxygen species (ROS) [55–56] and showed dominant expression in affected samples (Fig. 5B), *GPX1* and *GPX3* reduce ROS level to alleviate oxidative stress [57]. Moreover, expression of thymosin beta 4 (*TMSB4X*) is associated with phenotypic modulation of smooth muscle cells and angiogenesis [61–62], whose down-regulation contributes to developing larger atherosclerotic plaques in mice [61]. Since *TMSB4X* was also up-regulated in WB chickens [2 (unpublished data)], its particular role in veins and muscle remains elusive and worth studying in the future. Overall, these results suggest a possible role of veins in generating ox-LDL through endothelial and/or smooth muscle cell-induced oxidation of LDL, in turn leading to perivascular lipid deposition and LLM accumulation. The findings further support the previous hypothesis that venous endothelial cells in WB exhibit an increased level of metabolic activity and potential dysfunction [64].

### Myositis

Lysosomes are the principal organelle in charge of protein turnover and tissue homeostasis, involving in various pathways such as metabolism, autophagy, and so on [26]. In alignment with functional analysis (Fig. S2C), overexpression of multiple cathepsins (*CTSA, CTSB, CTSC*, *CTSD*, *CTSK, CTSS, CTSZ*) and lysosomal membrane proteins (*LAMP1, LAPTM5*) in the myositis cluster strongly indicated active lysosomal function, suggesting a dynamic turnover of damaged or inflamed myofibers [26–27]. Specifically, *CTSB* hydrolyses various myofibrillar proteins, including myosin heavy chain, tropomyosin, troponin T and I [26]. The co-expression of *CTSB* and its inhibitor cystatin-A (*CSTA*) further implied localized inflammation, as observed in pancreatic tumor cells [28]. The existence of myofiber inflammation was also supported by expression of CD74 molecule (*CD74*), MHC class II beta chain 2 (*BLB2*) and ferritin heavy chain 1 (*FTH1*). As the receptor for macrophage migration inhibitory factor, *CD74* regulates immune response as well as cell proliferation, and is implicated in other lipid-induced inflammatory diseases such as atherosclerosis [32]. Similarly, *BLB2* is related to adaptive immune response in chickens through antigen processing and presentation [63]. And ferritin is a pro-inflammatory cytokine whose expression matches inflammatory activity and remains up-regulated during muscle regeneration [33–34]. Furthermore, ferritin was also up-regulated in response to muscle damage attributed to reactive oxygen species after denervation [54], which was indicative of the oxidative stress related myofiber damage in WB.

Considering the high level of oxLDL and ubiquity of oxidative stress in WB [1–2], lysosomal membrane permeabilization (LMP) could be induced and result in leakage of cathepsins and lysosome-mediated apoptosis [29]. Moreover, this muscle inflammation was supported by *CTSS* and *CTSZ*, whose expression compensates for the activity of *CTSB* and *CTSC* during LMP-regulated inflammation [29]. Such crosstalk between lysosome and cell death was also observed in bovine muscle postmortem, where ROS level was positively correlated with lysosomal membrane stability and activities of *CTSB* and *CTSD* [30]. Although little is known about the biological function of IFI30 lysosomal thiol reductase (*IFI30*) in muscle, *IF130* was believed to dampen oxidative stress by recycling oxidized glutathione in hematopoietic stem cells [31]. That said, this spatial transcriptomic analysis displayed a possible linkage between lysosome-mediated muscle degeneration and inflammation probably under oxidative stress in WB, which warrants future study in a more advanced stage of this disease using molecular approaches.

### Muscle heterogeneity

As stated by pathway analysis, myofiber cluster M2 manifested a normal expression pattern in accord with type II muscle fueled by glucose typical for avian pectoral muscle. Even though no pathologic lesions were observed in any of the histological images other than myositis, myofiber cluster M3 and M4 exhibited a profile consistent with WB denoted by markers identified in previous studies.

Majority of the top marker genes of the myofiber clusters M3 and M4 clusters were ubiquitously found up-regulated in the affected muscle samples and are involved in myogenesis, cell proliferation and regeneration [3,43–45]. For instance, expression level of myogenin (*MYOG*) and myosin heavy chain 15 (*MYH15*) increased linearly with WB severity to participate in skeletal muscle repair [43]. Cysteine and glycine rich protein 3 (*CSRP3*) and its grouped protein myozenin 2 (*MYOZ2*) are related to repair mechanism and fiber-type switching in WB affected muscle [3,44]. Musculoskeletal, embryonic nuclear protein 1 (*MUSTN1*), ankyrin repeat domain 1 (*ANKRD1*), HOP homeobox (*HOPX*) and PDZ and LIM domain 3 (*PDLIM3*) were up-regulated in WB birds at 7 weeks [45]. Particularly, *MUSTN1* showed differential expression between slow growing chickens and those affected with WB at 6 weeks of age [67]. Their up-regulation in affected samples at both 6-7 weeks and 23 days post hatch implied a constant myogenesis and regeneration process in broiler breast muscle probably due to the fast muscle growth. Meanwhile, the resulted muscle hypertrophy should be accompanied with cytoskeleton assembly in order to maintain a structural organization and contractibility [46]. That being said, a gene in charge of cytoskeleton reorganization, muscle RAS oncogene homolog (*MRAS*) was also up-regulated in WB affected chickens of 7 weeks [5]. Overall, the results of this study reaffirm the importance of these WB-specific markers and identify their heightened expression specifically within myofibers.

### Conclusions

Via spatial transcriptomics, distinct histologic features were successfully characterized based on their gene expression profiles, including LLM, myofibers, myositis, connective tissue and vasculature. In our prior bulk RNA-seq studies, we observed increased expression of lipid metabolism genes during the early stages of WB [45, 65–66]. However, the specific cell type and regions responsible for this gene expression signature remained largely unknown. Our results from spatial transcriptomics revealed LLM as the key cell population responsible for altered lipid metabolism during early stages of WB. Building upon prior findings of elevated 9-HODE and 13-HODE in WB [4], we propose that the formation of LLM in broiler chickens occurs as a cellular cascade stimulated by oxLDLs. Additionally, we report discrepancies in gene expression within connective tissue and vascular regions, leading to discoveries of potential WB specific biomarkers and generating insights in muscular disease pathology. We also confirmed existence of gene expression heterogeneity within chicken pectoralis major muscle, despite visually indistinguishable histological presentation across intact myofibers. In inflamed myofibers, active participation by lysosomes in muscle homeostasis was suggested by the up-regulation of numerous lysosomal genes. Overall, this study revealed potential gene expression biomarkers related not only to myopathic muscle but also to various cell types present in healthy pectoralis major muscle of broiler chickens.

## Materials and Methods

### Chickens and tissue collection

As part of an ongoing genetic study of WB, 114 Cobb500 broiler chickens were raised in a chicken house at the University of Delaware and allowed free access to feed and water. The current experiment utilized p. major muscle tissues from three randomly-sampled chickens at 23 days of age, harvested post-euthanasia, and examined for gross lesions during necropsy. Specifically, chickens were examined for muscle firmness by manual palpation of the p. major muscle, and each bird was given a wooden breast score, as described in a previous work from our laboratory [24]. Approximately, 1 cm^3^ muscle samples were dissected from the cranial part of the right pectoralis major muscle and immediately embedded in Optimal Cutting Temperature (OCT) compound media, flash frozen in a bath of isopentane and liquid nitrogen before being stored at −80°C until further processing. The animal conditions and experimental procedures used in this scientific study were approved by the University of Delaware Institutional Animal Care and Use Committee (protocol number: 120R-2021-AP).

### Permeabilization experiments

The fresh frozen samples in OCT media were sectioned to 10-µm using a cryostat and mounted on 10X Visium slides. Tissue permeabilization (TP) experiments were conducted on both longitudinal- and cross-sectional muscle tissues as per manufacturer instructions using 10x Genomics Visium Tissue Optimization Kit (PN-1000193), yielding optimal TP time of 18 and 6 minutes, respectively. In the subsequent steps (section below), since samples 1 and 2 were largely longitudinal; TP time for this samples was set at 18 minutes during spatial RNA sequencing library preparation (i.e., the following section). However, for sample 3, which was an oblique section, the entire spatial transcriptomic experiment was replicated with 6 and 18 minutes TP time, mainly because the orientation of individual myofibers (obliquity) varied considerably (from cross-sectional to longitudinal) across different areas within the same tissue section. Therefore, repeating the experiment under optimal TP time for obliquity affected sample allowed capture of as much useful information as possible.

### Spatial RNA sequencing library preparation

The fresh frozen samples in OCT media were sectioned to 10-µm using a cryostat and mounted on 10X Visium slides and stained with hematoxylin and eosin (H&E). The H&E stained samples underwent high-resolution imaging on the confocal Dragonfly Spinning Disk and Super Resolution Microscope at the Bioimaging center of Delaware Biotechnology Institute (DBI) and were subsequently visualized using Fiji software [8]. After imaging, muscle samples were permeabilized and cDNA libraries were generated following the manufacturer’s instructions using Visium Spatial Gene Expression Slide & Reagent Kit (10x Genomics, PN-1000187; Pleasanton, CA). Quality analysis of the extracted total RNA and cDNA was conducted using the fragment analyzer to verify both quality and quantity, including Qubit quantification. The libraries were then sequenced in paired-end 150 bp mode on an Illumina Nextseq 2000 system (Illumina, San Diego,CA), with sequencing depth determined by the percentage capture areas covered by each muscle sample.

### Histopathologic analysis and tile annotation

Microscopic analysis and annotation were conducted on a tile-by-tile basis for each image file captured per specimen by co-author P. Khondowe. The classification of samples into WB affected and unaffected categories was based on histopathologic features identified by co-author E. Brannick, an ACVP certified veterinary anatomic pathologist as described previously [24]. Specifically, tiles were annotated for evidence of perivascular lipid-laden macrophages (LLM), inflammatory infiltrates within the interstitium or myofibers (myositis) or myofiber degeneration typical of WB. Structurally, regions of connective tissue, blood vessels, and white adipose were also labeled. Once all tiles were annotated, the veterinary pathologist (co-author E. Brannick) reviewed all tiles/annotations for accuracy and compiled data regarding the microscopic lesions and architectural elements present in each tile for subsequent data analysis and interpretation with regional molecular expression profiles as described below.

### Statistical analysis

The fiducial frames were manually aligned for each slide using Loop Browser v6.2.0 software (10xGenomics). Subsequently, sequencing data were aligned to the chicken reference genome Gallus_gallus-6a (Ensembl, database version 99) and overlaid onto the histological images using the SpaceRanger v2.0.0 (10xGenomics) to quantify gene expression and obtain the feature-by-barcode expression matrix. Unbiased k-means clustering was performed using Loup Browser v6.2.0 to determine feature clusters with k set as 10 to increase the power of cell type differentiation. Marker genes for each identified cluster were selected as up-regulated genes (adjusted P-value < 0.1).

Integration analysis was conducted using Seurat v4.3.0 [9] in R v4.1.0 [10], in a way that all datasets were anchored together to align the same cell types across conditions. Briefly, the 2000 most variant genes were identified as shared sources of variation across all the samples were identified by the canonical correlation analysis [11]. Before integration, anchors between datasets were extracted based on gene expression values followed by similarity score correction to remove incorrect anchor pairs [11]. This time cell type clusters were generated by the unsupervised shared nearest neighbor (SNN) graph-based clustering with resolution parameter set at 0.5 as it yielded clusters with meaningful biological interpretation without over-clustering the data [11,12]. Uniform Manifold Approximation and Projection (UMAP) was constructed to visualize integrated data and clusters using Seurat v4.3.0 [9] in R v4.1.0 [10]. Marker genes were extracted for each cluster using default Wilcoxon Rank Sum test in Seurat v4.3.0 [9] to select only up-regulated genes (adjusted P-value < 0.05) expressed in at least 25% of Visium spots in a cluster compared to all other spots. Pathway enrichment analysis was conducted using the ToppFun function of ToppGene Suite, with a false discovery rate threshold of 0.05 [13].

## Acknowledgements

This research is supported by U.S. Department of Agriculture, Agriculture, and Food Research Initiative competitive grant number 2021-67015-34543. We would like to thank Jeffery Caplan, Jean Ross, and staff from the University of Delaware Bio-imaging center; along with Bruce Kingham and Mark Shaw from University of Delaware Sequencing and Genotyping Center at the Delaware Biotechnology Institute for their expertise and assistance with sample processing, imaging and RNA sequencing. We also thank Dr. Carl Schmidt for his insightful input during manuscript preparation.

## Author Contributions

B.A. conceived of the study, designed, and planned the experiments, procured funding support, provided advisory guidance, revised and edited the manuscript. E.B. conducted tile and histology analysis, revised and edited the manuscript. P.K. performed data curation, tile analysis, revised and edited the manuscript. Z.W. performed data curation, analysis and wrote the manuscript.

